# Deciphering the unique SNPs among leading Indian Tomato Cultivars using Double Digestion Restriction Associated DNA sequencing

**DOI:** 10.1101/541227

**Authors:** Reddaiah Bodanapu, Sreehari V. Vasudevan, Sivarama Prasad Lekkala, Navajeet Chakravartty, Krishna Lalam, Boney Kuriakose, Chetan Bhanot, Pallavi Namdeo Morey, A. V. S. Krishna Mohan Katta, George Thomas, Saurabh Gupta, V B Reddy Lachagari

## Abstract

World-wide grown and consumed tomato (*Solanum lycopersicum*) crop used as model system for new cultivar and fruit development. Genetic and genomic research of Indian tomato cultivars will provide an insight to develop new breeding strategies and crop improvement. The present study, aimed to identify the high quality common and unique single nucleotide polymorphisms (SNPs), present in 9 different Indian tomato cultivars using double digestion restriction associated DNA sequencing (ddRAD-seq). Total of 36,847,092 raw reads (3.68 GB) were generated for all samples and 3,329,625 of high-quality reads were aligned uniquely to the reference tomato genome. Using stringent filtering, a total of 1,165 SNPs and 69 INDELs were found in genic regions, along with the unique variants to each cultivar was observed. Similarly, 7 and 33 variants were identified in chloroplast and mitochondrial genome of tomato. In addition, the population structure and genetic relationship among these cultivars suggested 4 well-differentiated sub-populations. Functional annotation of SNP/INDLEs associated with flanking sequences along with gene ontology and pathway analysis was performed. Identified SNPs/INDELs could be useful as markers for variety identification for genetic purity analysis. Findings from this work will be useful to plant breeders and research community to deepen their understanding and enhance tomato breeding programs.

## Introduction

Tomato (*Solanum lycopersicum*) is a highly consumed vegetable across the world; originated from South and Central America and spread to the rest of the world with accompanying morphological diversification. India is the second largest producer of tomato after China. Notably, the fruit colours, sizes, shapes, tastes and flavours of various cultivars are being associated with local environments and gastronomies (Nag 2017, Khan 2017). Among the Indian states, Andhra Pradesh holds the top position in terms of production of tomato (Monthly Report Tomato January 2018). Varieties such as Pusa, Pusa-120, Ruby, HS-101, HS-102, Hisar Arun, and Hisar Lalit are endorsed for cultivation across India. In addition, numerous other cultivars i.e. Kashi Aman, Kashi Abhiman, Arka Ananya, Arka Vardhan, Arka Saurabh, Arka Meghali, Arka Vishal, Arka Abhijit, Arka Ahuti (Sel-11) Arka Vikas (Sel-22), Arka Abha (BWR-1), Arka Alok (BWR-5), Kashi Amrit, Kashi Anupam, Kashi Sharad, Pusa Sheetal, Pusa Gaurav, Pusa Rohini, Pusa Hybrid-2, Pusa Hybrid-4 and Pusa Hybrid-8 are widely used for cultivation in different part of India (Major Singh 2016). Genome sequencing of 84 tomato accessions which includes wild species of *S. lycopersicon, S. arcanum, S. eriopersicon* and *S. neolycopersicon* have been conducted under 100 Tomato genome projects. Recently,150 tomato genome re-sequencing project was established and started to explore in-depth genetic variation available in tomato (Sato 2016, Aflitos 2014, Causse 2013, Lin 2014). These attempts were aimed to categories genome-wide Single Nucleotide Polymorphisms (SNPs) within *S. lycopersicum* along with shared polymorphisms among closely related species (Shirasawa 2013). Additionally, numerous interspecific genetic linkage maps have been constructed between well-known cultivars of tomato and were used to identify the responsible genes for interspecific and intraspecific phenotypic variations (Tanksley 2009, Lin 2014).

High throughput sequencing and genotyping methods have been playing a crucial role in the progress of genomics and genetics. Rapid progress in next-generation sequencing (NGS) technologies have made it easy generate a large number of SNPs in both model, non-model crop and vegetable plant species (Cloutier 2016, Davey 2011). Congruently, double digest restriction site-associated DNA sequencing (ddRAD-Seq) technology has recently become more popular technique due to its flexibility, low cost, and advantage over genotyping by sequencing (GBS) and restriction site-associated DNA sequencing (RAD-Seq) (Peterson 2012, Elshire 2011, Baird 2008). In ddRAD-Seq two restriction enzymes are employed for digestion of genomic DNA which reduces the time and cost to prepare the sequence libraries, allows paired-end sequencing (Peterson 2012). In present study, ddRAD-Seq was performed for 9 well known tomato cultivars such as Arka Abha (BWR-1), Arka Ahuti (Sel-11), Arka Alok, Arka Ashish, Arka Saurabh, Arka Meghali, Arka Vikas (Sel-22), Arka Vikas-vir and PKM1 to annotate and investigate the unique variants across aforementioned lines to identify individual markers. Subsequently, the effects of these variants on different gene function were also investigated and explored for marker assist selection and breeding of the tomato cultivars.

## Experimental Protocol

### Plant material and Genomic DNA isolation

All the Arka series of lines were obtained from IIHR, Bengaluru, India. PKM1 was obtained from TNAU, Coimbatore, Tamil Nadu, India. Nine tomato cultivars Arka Abha (BWR-1), Arka Ahuti (Sel-11), Arka Alok (BWR-5), Arka Ashish, Arka Meghali, Arka Saurabh, Arka Vikas (Sel-22), Arka Vikas_Vir and PKM1 were cultivated in the field of AgriGenome Labs Pvt. Ltd. at Hyderabad, India. 3 weeks old plants juvenile leaves were collected, and genomic DNA was isolated using the DNeasy Plant Mini Kit (Qiagen, Hilden, Germany), and quantitated using Eppendorf (BioSpectrometer fluorescence, Germany), Qubit fluorometer (Life Technologies, Carlsbad, CA, USA) and agarose gel electrophoresis.

### Selection of restriction enzymes and adapter design

*Mlucl* and *Sph1* enzyme pair is capable for producing higher number of fragments in a broad range of plant species (Burns 2017). Hence same enzyme combination was used to generate higher number of fragments of 9 Tomato cultivars. 1μg genomic DNA of each cultivar was double digested with SphI and *MlucI* restriction enzymes and incubated at 37°C for 16-20 hours. Cleaning the unwanted restricted fragments using AMpure XP beads (Beckman Coulter Genomics). Ligation of P1 (Barcoded) and P2 adapters was carried out using T4 DNA ligase. Cleaning the unwanted ligated fragments using AMpure XP beads (Beckman Coulter Genomics). Ligated product was fractionated in 2% SybrSafe agarose gel electrophoresis to check the product size in between 250 −350 bp (Peterson 2012).

### Library preparation and sequencing

Quality and quantity check of ligated sample was performed to increase the concentration of sequencing libraries, PCR amplification (8-12 cycles) was performed by adding Index 1 and Index 2 (8 nt long) for multiplexing using Phusion™ Taq DNA polymerase kit. Products of PCR amplification was analysed in Agilent bioanalyzer to quantify molarity and fragment size distribution (Peterson 2012). For each sample, individual libraries were prepared and included in one lane. The sequencing was done based on V4 chemistry on HiSeq4000 platform which uses different library index sequences.

### Pre-processing, mapping and variants annotation

The reads were filtered based on presence of specific rad tags, followed by base trimming and adapter trimming. The quality distribution plot was generated using FASTQ and filtered data was aligned to reference tomato genome along with its mitochondria and chloroplast genome obtained from Sol Genomics (vSL3.0) and NCBI respectively, using Bowtie2 (v2.2.2.9) (Langmead 2012). The variant calling was performed based on the alignment of the samples with the reference genome using SAMtools (v0.18) (Li 2009). The variants were filtered based on the read depth and high-quality variants were reported. In addition, polymorphic homozygous markers were identified between the cultivars using inhouse PERL scripts. Further, functional annotation of the identified variants associated genes was performed using SnpEff (v 3.1) (http://snpeff.sourceforge.net/).

### Zygosity, diversity, phylogenetic and kinship analysis

The zygosity analysis, diversity analysis, phylogenetic and kinship analysis, for 9 cultivars were carried out based on the genotype data at read depth 10 with MAF ≥ 0.05. The dendogram was constructed based on the genotype data using similarity matrix generated by NJ module. The degree of kinship among individuals were estimated based on the genotype using VanRaden’s method (VanRaden 2008). Kinship analysis was studied using Centered-IBS matrix using TASSEL5 (v5.2.28) (Bradbury 2007). Further, functional annotation of the genes associated with individual unique and common variants were identified.

## Results and discussion

### Samples collections, raw data analysis and alignments

Genome sample of each cultivar was digested with *Mluc1* and *Sph1* restriction digestion enzyme having different frequencies of recognition sites. Quality check of fragmented samples indicates that all qualified screening criteria. Furthermore, quality check, screening and filtering of raw data reveals different read statistics as depicted in Table 1. Highest and lowest number of reads with RAD Tag was found in Arka Abha (BWR-1), and Arka Vikas-Vir respectively, while Arka Ashish and Arka Saurabh encountered with highest and lowest number of reads with no RAD Tag respectively. The percentage of uniquely aligned reads from Arka Abha (BWR-1), Arka Ahuti (Sel-11), Arka Alok (BWR-5), Arka Ashish, Arka Meghali, Arka Saurabh, Arka Vikas (Sel-22), PKM1 & Arka Vikas -Vir with Sol Genomics SL v3.0 reference genome was 93.56, 94.38, 92.86, 94.19, 94.42, 93.22, 93.36, 93.30, & 94.72 respectively. Overall, 87.70% of reads were mapped to reference genome, out of them 93.77% of reads were mapped uniquely. On average, 3.97% and 4.69% of the reads from each sample were aligned to mitochondria and chloroplast genome respectively. Density plot of uniquely aligned reads with reference tomato genome (Figure 1) indicates their distribution over all 12 chromosomes of tomato.

**Table 1.** Sequanced reads measurements across selected 9 cultivar along with statistics genome-wide alignment and annotated SNPs & INDELs at read depth (RD) ≥ 10.

| Sample Names | # of reads with RAD-Tag in (R1+R2) | # of high-quality reads | # of aligned reads | % of aligned reads | % of uniquely aligned reads with nuclear genome | # of aligned reads with Mitochondria Genome | # of aligned reads with Chloroplast Genome | #SNPs at RD$\geq 10$ | Unique #SNPs at RD$\geq 10$ | #INDELs at RD$\geq 10$ |
| --- | --- | --- | --- | --- | --- | --- | --- | --- | --- | --- |
| Arka Abha (BWR-1) | 6,404,140 | 6,404,090 | 5,439,634 | 84.94 | 93.56 | 225,520 | 273,094 | 396 | 50 | 38 |
| Arka Ahuti (Sel-11) | 3,666,454 | 3,666,384 | 3,286,179 | 89.63 | 94.38 | 155,528 | 157,718 | 186 | 29 | 31 |
| Arka Alok (BWR-5) | 4,126,266 | 4,125,944 | 3,510,353 | 85.08 | 92.86 | 145,866 | 161,490 | 416 | 73 | 33 |
| Arka Ashish | 5,282,126 | 5,281,902 | 4,708,815 | 89.15 | 94.19 | 238,658 | 251,490 | 460 | 140 | 42 |
| Arka Meghali | 4,386,412 | 4,386,366 | 3,907,374 | 89.08 | 94.42 | 183,282 | 226,516 | 347 | 26 | 34 |
| Arka Saurabh | 4,529,000 | 4,528,704 | 3,809,998 | 84.13 | 93.22 | 156,462 | 182,570 | 349 | 21 | 35 |
| Arka Vikas (Sel-22) | 3,619,712 | 3,619,558 | 3,129,469 | 86.46 | 93.36 | 125,388 | 153,646 | 340 | 30 | 37 |
| PKM1 | 4,109,130 | 4,108,926 | 3,510,666 | 85.44 | 93.30 | 137,024 | 56,338 | 302 | 47 | 40 |
| Arka Vikas -Vir | 725,242 | 725,218 | 678,659 | 93.58 | 94.72 | 39,948 | 158,596 | 256 | 44 | 11 |

**Note:** # Indicates Number and % Percentage

**Figure 1:**
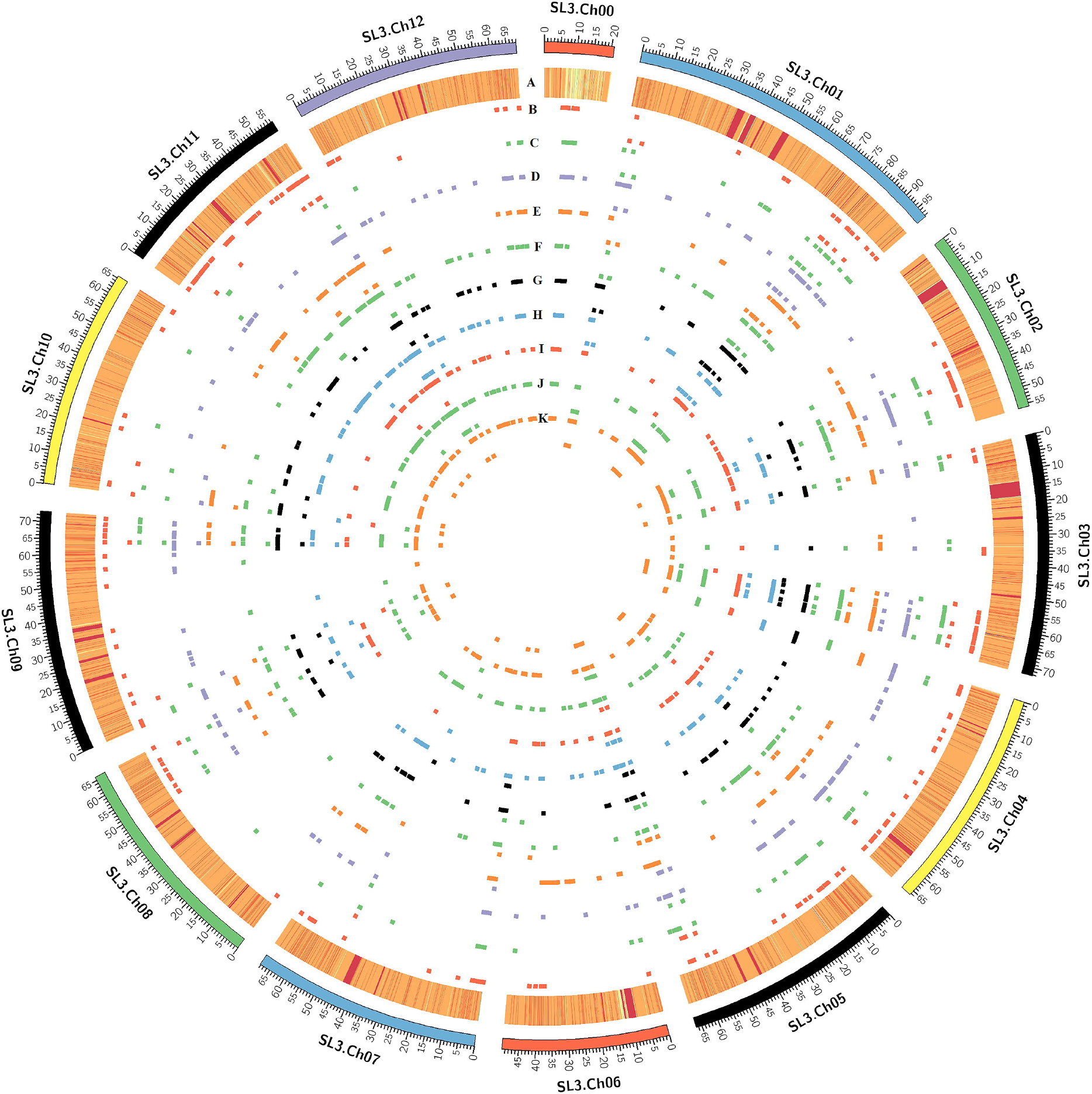
Chromosomal distribution of generated reads and identified SNPs & INDELs in each cultivar of tomato in a Circos plot. (A) number of generated base pair density in a chromosomal location shown heatmap. Distribution of identified SNPs(up) & INDELs(down) via scatter plots for all nine cultivars i.e. (B) Arka Abha (BWR-1), (C) Arka Ahuti (Sel-11), (D) Arka Alok (BWR-5), (E) Arka Ashish (F) Arka Meghali (G) Arka Saurabh, (H) Arka Vikas (Sel-22), (I) Arka Vikas-Vir and (J) PKM1.

### Identification of polymorphic variants and cultivar specific markers

A total of 3,353 high-quality SNPs/INDELs (read depth (RD) ≥ 10) were found to be present in genic regions across 9 cultivars when compared with nuclear genome (Table 1). The nuclear genome variants found in intergenic and genic regions were evenly distributed across all chromosomes of tomato genome (Figure1). In-depth analysis of genic region identified SNPs (1,169) and INDELs (69) distributed over all the chromosomes of tomato and wre considered for downstream analysis. Large number of SNPs in Arka Ashish (460) and Arka Alok (BWR-5) (416) while Arka Ahuti (185) share the lowest contribution (Supplementary Table S1). Considering the INDELs distribution among all cultivar indicate that Arka Ashish (43) share highest contribution (Supplementary tables S2). Further, unique SNPs among the 9 cultivars *viz.* Arka Abha (BWR-1) (50), Arka Ahuti (Sel-11) (29), Arka Alok (BWR-5) (73), Arka Ashish (140), Arka Meghali (26), Arka Saurabh (21), Arka Vikas (Sel-22) (30), PKM1(47) & Arka Vikas-Vir (44) were identified. Similarly, unique INDELs across all selected cultivars, only Arka Ahuti (Sel-11) (1), Arka Ashish (1), PKM1(2) & Arka Vikas-Vir (1) were reported (Supplementary Table S3). Annotation of SNPs were categorised into missense, 3’ 5’-UTR, splice region up & down stream gene variant, among them missense mutations which responsible for phenotypic trait change. Arka Abha (BWR-1) (19), Arka Ahuti (Sel-11) (6), Arka Alok (BWR-5) (17), Arka Ashish (22), Arka Meghali (14), Arka Saurabh (21), Arka Vikas (Sel-22) (17), PKM1(17) & Arka Vikas-Vir (18) along with other type of the SNPs variants are also identified (Supplementary Table S1).

### Identification of SNPs in chloroplast and mitochondrial genome

Total of 7 and 33 SNPs (RD≥ 10) were identified in chloroplast and mitochondrial genome respectively (Supplementary Table S4 & S4). Annotation of SNPs shows only one SNPs at 2,239 position in Mttb gene with variation A/T, found in cultivar Arka Meghali, PKM1 whereas A/G in Arka Saurabh. Mttb (Acc. No. NC_035963.1) is a SecY-independent transporter protein helps in proton motive force dependent protein transmembrane transporter activity.

### Genetic relationship analysis among tomato cultivars

This analysis predicts 6,957 homozygous polymorphic SNPs between 9 cultivars. Genotyping also mimics out 69,570 data points and 139,140 gametes based on genotyping data at read depth 10 with MAF cut off of 0.05. Zygosity analysis found 13,935 heterozygous calls with the 0.2003 proportion and it infers that Arka Ahuti (Sel-11) showed the highest percentage (78.63%) of heterozygosity followed by Arka Abha (BWR-1) (36.136%), Arka Alok (BWR-5) (31.594%) while PKM1 showed less heterozygosity indicating more homologues to reference (Figure 2A). Further, to investigate the genetic relationships between identified SNPs of all cultivar, a phylogenetic tree was constructed based on a pairwise distance matrix using NJ method (Figure 2B). The 9 cultivars with reference tomato genome were classified into 4 main clades on the basis of SNPs clustering: clade 1 consists of *S. lycopersicum* (3.00), Arka Ahuti (Sel-11), Arka Abha (BWR-1) and Arka Ashish; clade 2 consists of Arka Saurabh, Arka Vikas (Sel-22), PKM1; clade 3 having Arka Vikas-Vir, Arka Meghali and clade 4 only Arka Alok (BWR-5) showed higher divergence with others. The largest clade1, clade 2, contained two major subclades showing that they are the close orthologs. Similar analysis was also performed in onion inbred lines (Lee 2018) and grape cultivars (Laucou 2018).

**Figure 2:**
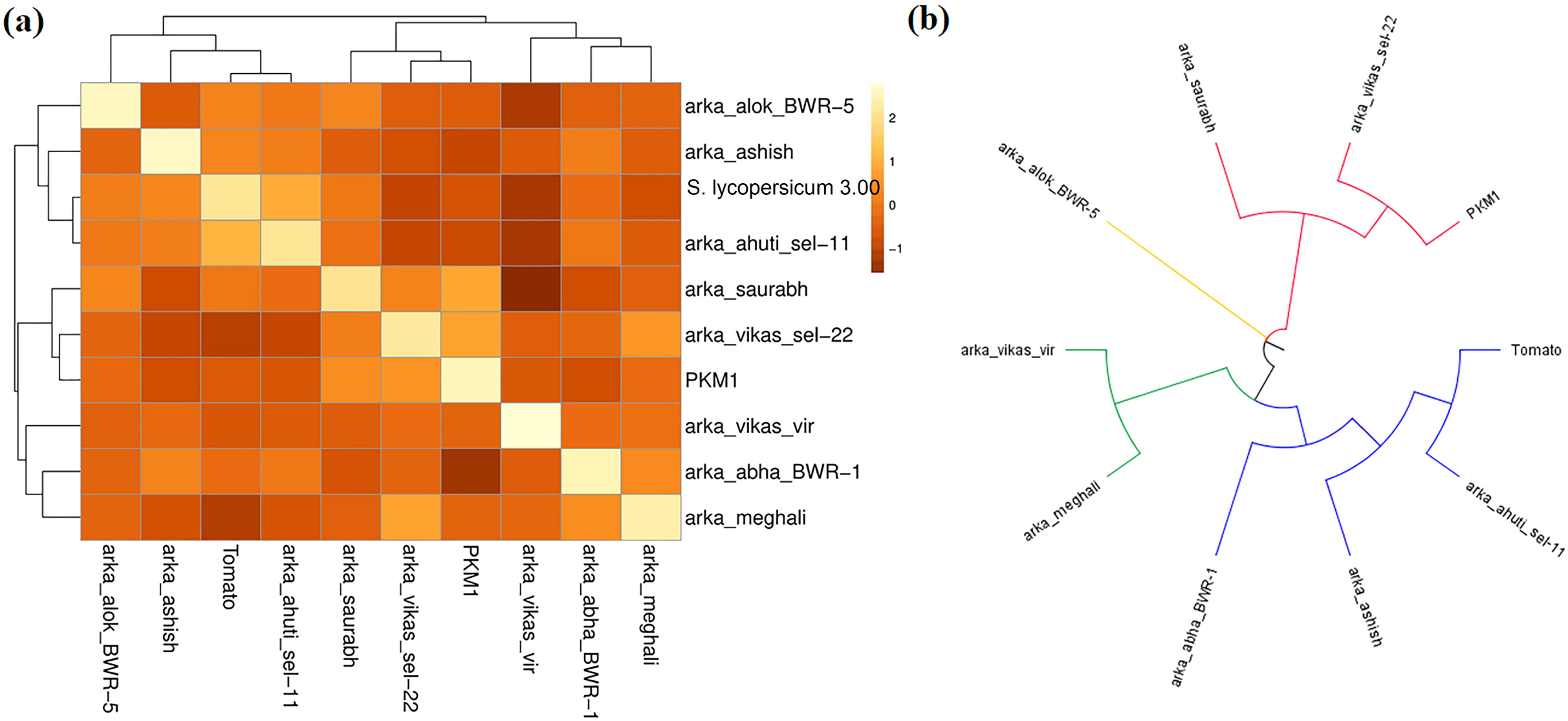
Genetic relationship analysis among all selected cultivars using reference S. lycopersicum 3.00 genome. (a) Heatmap of kinship analysis identified SNPs across all cultivar (b) linear tree diagram shows evolutionary divergence among the cultivars.

### Gene ontology examination of genic region SNPs

The gene ontology (GO) analysis of genic region SNPs was identified and GO terms belongs to biological process, molecular functions and cellular components shown in Figure 3. Transcription [GO:0006351] & regulation of transcription [GO:0006355] and ATP binding [GO:0005524] & Metal binding [GO:0046872] are highly enriched GO terms associated with biological processes and molecular functions respectively. Similarly, highly enriched GO terms associated with cellular components are integral components of membrane [GO:0016021], nucleus [GO:0005634]. GO terms associated with nutrient transportation, stress tolerance and resistance and other morphological traits of tomato (Gupta 2017, Gupta 2018a, Gupta 2018b)

**Figure 3:**
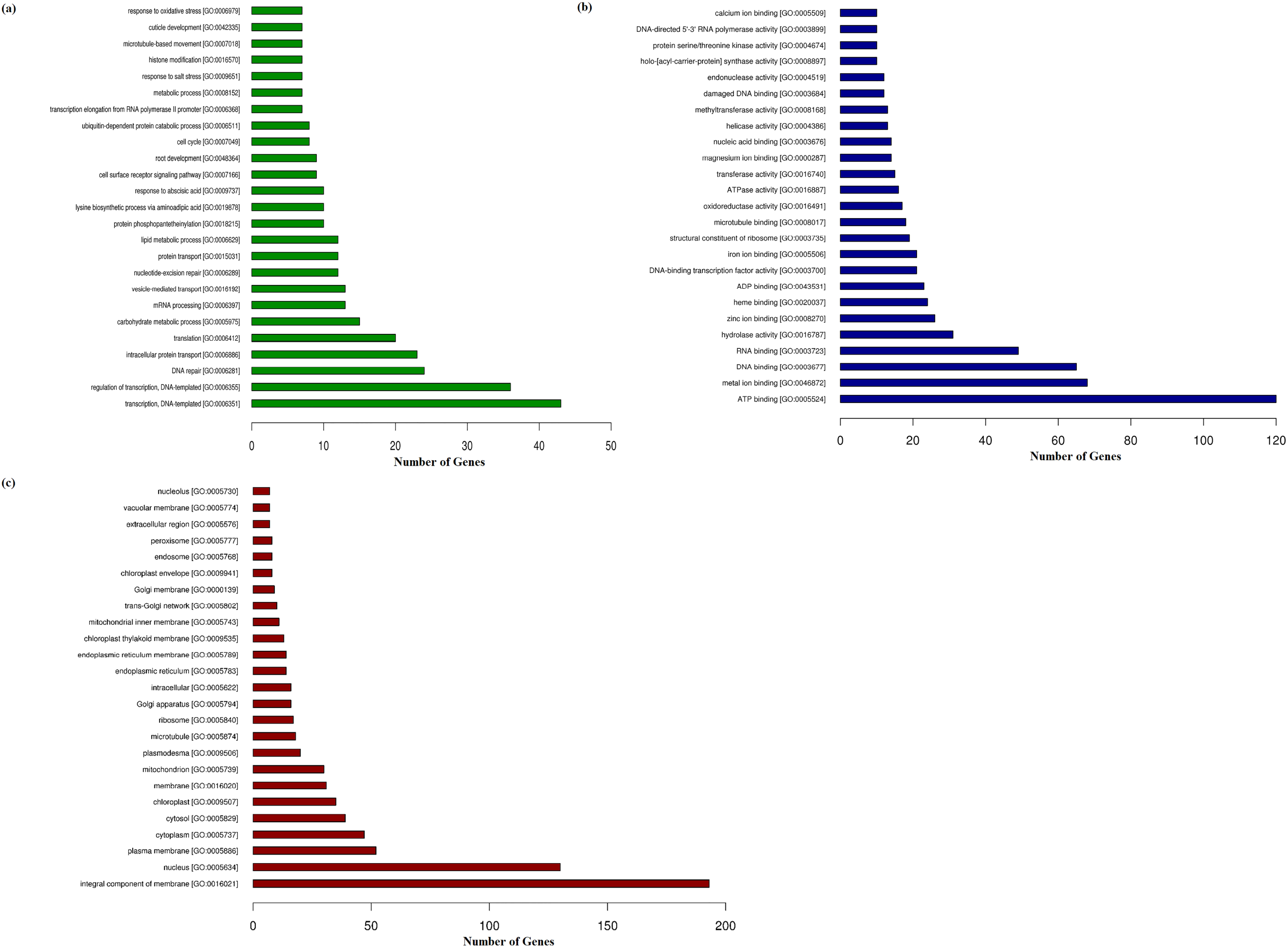
Gene ontology terms associated with the SNPs found in gene region of all cultivar. (a) GO terms associated with biological process (b) GO terms associated with Molecular functions and (c) GO terms associated with cellular components.

### Functional annotation of missense SNPs associated genes

The missense SNPs associated with the genes were identified (Supplementary Table S1&S2). Along with unique SNPs to each cultivar with their effects was studied (Figure 4). Seven missense SNPs of Arka Ahuti (Sel-11) associated with 7 genes in which 3 SNPs is unique for this cultivar. Auxin transport protein BIG, TCP transcription factor 4 and Gibberellin receptor GID1A have unique mutation K2704T, L515S and W289C respectively, possibly these mutations, enhancing the essential hormones that regulates growth and development (Murase 2008, Gil 2001). A total of 17 and 18 SNPs were found in Arka Vikas (Sel-22) and Arka Vikas-Vir, sharing some common SNPs, while they also have 1 and 3unique SNPs respectively. Three unique SNPs of Arka Vikas-Vir associated with 30S ribosomal protein S14 (T22A & R8H), and DNA repair endonuclease UVH1(G866S), while in case of Arka Vikas (Sel-22) gene is not functionally annotated. Twenty-one missense SNPs were observed in Arka Saurabh cultivar in which 3 SNPs are unique to this variety. These mutations found in Serine/threonine-protein kinase Nek1(His199N), Nucleotidyl transferase family protein (H322R), Programmed cell death 7(S313N), Cation/H (+) antiporter (N620D), RNA polymerase-associated protein RTF1(K185N), Dentin sialophospho protein-related (Q428Pro), DnaJ (A1224S), VHS domain-containing protein (G661D), Exostosin family protein (Phe307S), BTB/POZ domain-containing protein (D245E), Transducin/WD40 repeat-like superfamily protein (R672C), Intracellular protein transporter USO1-like protein (R240W), ARM repeat superfamily protein (N770S), Calcium-dependent lipid-binding-like protein (V2933Phe), Poly(A) RNA polymerase cid14(R1188His), Tetratricopeptide repeat (TPR)-like superfamily protein (V71D), Pentatricopeptide repeat-containing protein (L238V) and Pleiotropic drug resistance ABC transporter (A515Pro), may be responsible for various stress tolerance and resistance phenotypic traits of this cultivar (Gupta 2018b, Gupta 2017b). Out of 19 missense SNPs, 3 unique SNPs are found in Arka Abha (BWR-1) and mutating the disease resistance protein RPP5(D22E), Alpha/beta-Hydrolases superfamily protein (V306Ile) and Proteasome-associated ECM29-like protein (A784D) enhancing the disease tolerance, aligning with this cultivar characteristics (Belkhadir 2004). Eight unique missense SNPs are reported out of 22 SNPs in Arka Ashish and are associated with Cell division cycle protein 27(K575T), N-glycosylase/DNA lyas (R16His), Carbohydrate-binding-like fold protein (K1538R), Plant cadmium resistance 10 (S9L), Zinc finger C2H2 type (T233A) and Histidine-tRNA ligase (T423P). Combined effect these SNPs possibly participating in physiological traits and tolerant to powdery mildew characteristic. Arka Alok (BWR-5) is bacterial wilt resistance variety with medium size fruit having 17 SNPs in which 3 unique SNPs directing mutations in Pntatricopeptide repeat-containing protein (E498G), Mitochondrial trehalose-6-phosphate synthase-6 (V466A) and GI protein (A1011P). Plant specific nuclear GI protein involved in variety functions *viz*. flowering time regulation, light signalling, herbicide tolerance, cold tolerance, drought tolerance, hypocotyl elongation, control of circadian rhythm, sucrose signalling, starch accumulation, chlorophyll accumulation, transpiration, and *miRNA* processing (Mishra 2015, Gupta 2019). Arka Meghali consist of 14 missense SNPs associated with Serine/threonine-protein kinase Nek1, Nucleotidyl transferase family protein, Programmed cell death 7 protein, Dentin sialophosphoprotein, VHS domain-containing protein, BTB/POZ domain containing protein, Intracellular protein transporter USO1-like protein, ARM repeat superfamily protein, Bidirectional sugar transporter SWEET, Peroxidase, Dynein-1-alpha heavy chain, flagellar inner arm I1 complex and ARM repeat superfamily proteins having mutation of H199N, H322R, S313N, Q428P, G661D, D245E, K232I, N770S, N70K & P45A, K91M, A6V & F7C and T510A respectively, out these no missense SNPs is unique for this cultivar. Besides Arka varieties, we also identified 17 SNPs in PKM1 cultivar of these 4 SNPs are unique and forming the mutations in proteins Brefeldin A-inhibited guanine nucleotide-exchange protein 2 (K876R), Mediator of RNA polymerase II transcription (T2N), Microtubule-associated protein (A147P) and Gibberellin receptor GID1A (G301C) possibly control the regulation of transcription and nutrient transportation (Murase 2008, Blazek 2005).

**Figure 4:**
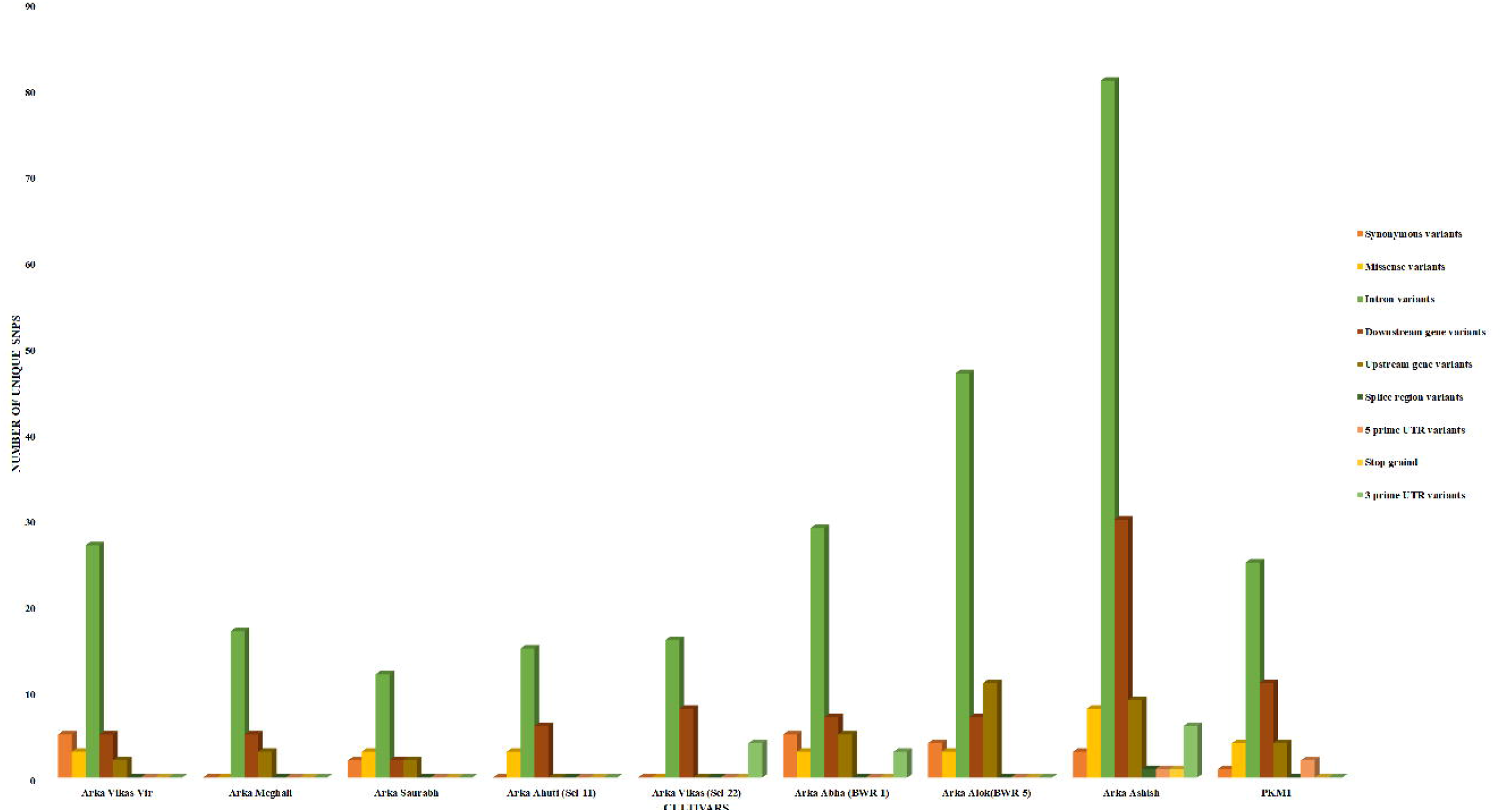
Cultivar-wise distribution of identified unique SNPs with their effects.

### Functional annotation of INDELs containing genes

The missense INDELs, associated with the genes were identified. The INDELs that are unique to one cultivar were identified (supplementary Table S2). Highest INDELs were found in Arka Ashish (42) among all Arka cultivar, followed by Arka Abha (BWR-1) (38), Arka Vikas (Sel-22) (37), Arka Saurabh (35), Arka Meghali (34), Arka Ahuti (Sel-11) (30). Arka Ashish share only 1 unique insertion of ‘TA’ in place of ‘T’ in the gene encoding kinase family with ARM repeat domain-containing protein. Similarly, we also observed unique deletion ‘CTTTTTTTTTT’ in gene encoding Importin subunit beta-1 protein and ‘CAAAAAAAAA’ in gene encoding Ubiquitin-like-specific protease ESD4 protein of Arka Ahuti (Sel-11), and Arka Vikas-Vir respectively. PKM1 showed 40 INDELs in which most of them are like the Arka cultivar only 2 INDELs found unique to this cultivar. Identified unique and common deletions could be helpful for marker assist selection.

## Conclusions

We identified cultivar specific unique as well as common SNPs and INDELs for leading tomato cultivars of India using ddRAD-seq analysis. Identified SNPs with their genetic characteristic and functional annotations, that will be useful for designing the markers for variety identification through genetic purity analysis. Besides, identified information can be useful for plant breeders to develop new tomato cultivar.

## Acknowledgments

Authors would like to acknowledge to ICAR-Indian Institute of Horticulture Research for providing the Arka series tomato seed samples and Tamil Nadu Agricultural University for providing PMK1 tomato seeds.

## Contributions

BR, VBRL, SPL, KM, GT, SM, SG conceived the work and designed the experiments. SVV, LK, NC, BR, SG performed *in silico* and *in vitro* experiments.SG, VBRL, NC, SPL, BR analyzed the results. SG, VBRL, SPL, GT, CB, PNM contributed to writing the manuscript and discussed the results and commented on the manuscript.

## Compliance with Ethical Standards

This research did not involve any experiment on humans or animals. Data generated in this work was related to rice plant and submitted in SRA database having BioProject ID: PRJNA484084. Hence, all the authors declare that there is no non-compliance with ethical standards.

## Supplementary Tables Information

**Supplementary Tables S1:** Identified SNPs along with its annotation in (A) Arka Vikas-Vir (Sel-22), (B) Arka Vikas, (C) Arka Ahuti (Sel-11), (D) Arka Meghali, (E) Arka Saurabh, (F) Arka Abha (BWR-1), (G) PKM1, (H) Arka Alok (BWR-5) and (I) Arka Ashish tomato cultivars.

**Supplementary Tables S2:** Identified unique SNPs in cultivar (A) Arka Vikas-Vir, (B) Arka Vikas (Sel-22), (C) Arka Ahuti (Sel-11), (D) Arka Meghali, (E) Arka Saurabh, (F) Arka Abha (BWR-1), (G) PKM1, (H) Arka Alok (BWR-5) and (I) Arka Ashish tomato cultivars.

**Supplementary Table S3:** Identified total INDELs in cultivar (A) Arka Ahuti(Sel-11), (B) Arka Vikas-Vir (C) Arka Saurabh (D) Arka Vikas (Sel-22), (E) Arka Abha (BWR-1), (F) Arka Alok (BWR-5) (G) Arka Ashish (H) Arka Meghali, and (I) PKM1 Arka Ashish, in each sheet the yellow highlighted colour shows unique INDELs to that tomato cultivars.

**Supplementary Table S4:** SNPs identified in chloroplast genome across all tomato cultivars

**Supplementary Table S5:** SNPs identified in mitochondria genome across all tomato cultivars.

